# Life Histories and Study Duration matter less than Prior Knowledge of Vital Rates to Inverse Integral Projection Models

**DOI:** 10.1101/2024.04.06.588423

**Authors:** Connor D. Bernard, Michael B. Bonsall, Roberto Salguero-Gómez

## Abstract

1. Ecology has been surprisingly slow to address the uncertainty and bias that results from using short-term time series to draw long-term inference. To improve our understanding of assumptions around the temporal structure of vital rates (*e.g.*, survival, reproduction), we need tools that are feasible and capture longer-term, state-structured population dynamics.
2. Here, we use inverse modelling of a set of integral projection models (IPMs) to show how demographic rates can be accurately reconstructed from state-structure fluctuations in a population time-series. We use a particle-filtering optimisation algorithm to fit vital rates from time-series of varying length, parameter combinations, priors, and life histories.
3. We show how key life history traits such as generation time have little effect on the ability of our approach to accurately identify vital rates using state structure over time. Further, contrary to our expectations, the duration of our time-series data has relatively modest impact on the estimation of vital rates compared to the critical role of prior knowledge on vital rates.
4. ur framework to estimate IPM vital rates highlights the potential of inverse models to extend time-series for demographic models, but also demonstrates that long-term time-series are not a perfect surrogate for detailed demographic inference. We discuss the need for more work exploring the conditions when inverse modelling is an adequate tool based on species traits.

## Introduction

Ecology has been surprisingly slow to address the uncertainty and bias that results from using short-term time series to draw long-term inference (Shoemaker et al. 2013). Inferences of transient (short-term) dynamics (Hastings & Higgins 1994) and asymptotic (long-term) dynamics (Caswell 2002) in population ecology are only as robust as the data from which they are estimated. Sampling of vital rates (*e.g.,* survival, growth, reproduction) is often poor with respect to species with long generation times (see, for example, lifespan versus mean age of populations, Conde et al. 2019). Thus, in the absence of long-term studies that can adequately capture the full life cycle of long-lived species, short-term time series have served as a basis for many demographic studies and population projections (but see Clutton-Brock & Sheldon 2010; Gaillard, Coulson, & Festa-Bianchet 2010; Reinke, Miller, & Janzen 2019). Indeed, 40% of the >11,000 population models in the COMPADRE and COMADRE databases (Salguero-Gómez et al. 2015, 2016) and 24% of the 280 models in PADRINO (Levin et al. 2022) have been parameterised with ≤ 5 years of field data. This picture contrasts with the much longer generation times of the plants (median = 19.87 years (95% gamma C.I.: 3.36, 259.04; α = 0.301, β = 180.57); COMPADRE v. 6.22.5.0) and animals (median = 6.37 years (95% gamma C.I.: 2.15, 46.76; α = 0.42, β = 26.86; COMADRE (v. 4.21.8.0). Temporal replication of demographic studies is limiting even in databases with a clear focus on long-term ecology. For instance, only 60% of the studies in The Living Planet Index database exceed 10 years, while nearly 80% of records contain coverage <20 years (Living Planet Index 2022). Temporal limitations are also common across the wealth of long-term distributed monitoring networks, not limited to the Forest Global Earth Observatory (ForestGEO; Anderson-Teixeira et al. 2014), the Long-Term Ecological Research Network (I/LTER Callahan 1984; Nottrott et al. 1994), and the National Ecological Observatory Network (NEON; Schimel et al. 2007). In these cases, monitoring efforts are but a few decades old, in contrast to the typically high generation times of the species they aim to study (White 2019; Wolfe et al. 1987; Lindenmeyer et al. 2012; ForestGEO 2023).

Most estimators of population performance are agnostic to the specific length of the supportive time series data, such as the abovementioned time-series data. Indeed, asymptotic rates (*e.g.,* population growth rate, stable st/age structure), transient indices (*e.g.,* reactivity, recovery time; Stott 2011; Hastings & Higgins 1994), stochastic approximations of asymptotic rates (including Tuljapurkar’s small noise approximation (1982)), elasticities of population growth rate to vital rates like survival or reproduction (de Kroon et al. 1986), demographic buffering indices (Capdevila et al. 2020), and population viability analyses (Beissinger & Westphal 1998) can all be evaluated from long time series, but also in principle from as few as three time points for stochastic models or even two for deterministic models (Crone et al. 2011, 2013; Bierzychudek 1999). Thus, the expected stronger inference that long-term datasets confer to population forecasts is not communicated through the established metrics by which we currently characterise populations (Doak, Gross, & Morris 2005). Consequently, estimating near-term (*i.e.,* transient dynamics) and long-term performance (*i.e.,* asymptotic dynamics) from vital rates rests on assumptions about sampling density and temporal trends that might not be supported by underlying data (Pennekamp et al. 2019; Clutton-Brock & Coulson 2002).

Not all demographic methods are equally accessible or feasible. As such, compromises in the type of data sampled for demographic modelling may need to balance competing needs or practical limitations. Some data types are more commonly measured than others and may therefore be more frequently relied upon for demographic inference. The characteristic time-series duration for a given method, as well as its spread, may differ substantially. Examples include the overall population size and structure (*e.g.,* age-or stage-ratios) *vs.* individually tracked organisms from birth to death (*e.g.,* demographic surveys based on capture-mark-recapture methods (Lebreton et al. 1997)). Tracking individual vital rates can be challenging in remote locations or for species with cryptic stages (Paniw et al. 2017; Nguyen et al. 2019), may be difficult and/or harmful for certain species (Murray & Fuller 2000; Geen, Robinson, & Baillie 2019), or can require durations beyond doctoral studies, research grant tenures, and even researchers’ active careers in some instances. For these species, the only demographic data available may be those inferable from time-series population size and structurre.

Inverse models infer unobserved processes from observational data on the performance of a system (Keller 1976; Kaipio & Somersalo 2005). This family of models has a long history across scientific disciplines, including engineering (Keller & Cebeci 1972), physics (Glasko 1984), geology (Gottlieb & DuChateau 2012), and ecology (Gellner, McCann, & Hastings 2023). In contrast to a “cause-and-effect” approach, the inverse problem is “effect-and-cause” oriented, meaning that one uses lower dimensional realisations to infer higher order generating functions. Specific to population ecology, inverse problem approaches use population structure (*i.e.*, frequency of individuals in different st/age classes), abundances, and/or density to estimate unobservable vital rates. This approach uses *inverse models* in the sense of solving inverse problems (Wood & Nisbet 1991; Manly 1990; Caswell 2001; González et al 2016). Integrated population models (Schaub & Kerry 2021) are one of several methods to marshall different types of data to get at vital rate inference indirectly and often engage inverse model inference (Portillo-Tzompa, Martín-Cornejo, & Gonzales 2023).

Inverse models have gained renewed attention in ecology in recent decades. This focus has been facilitated by methodological advances in stage-structured projection models such as Integral Projection Models (IPMs) (Easterling et al. 2000), computational efficiency (algorithmic optimisation and Monte Carlo integration; Knape & De Valpine 2012), and accessibility of Bayesian approaches (Hamiltonian, Gibbs, Metropolist Hastings, and Sequential Monte Carlo algorithms; Scranton, Knape, & De Valpine 2014). Indeed, recent innovations have extended the potential utility of inverse models by relaxing assumptions regarding the determinism of the system (Ghosh, Gelfand, & Clark 2012) and with respect to stationarity of underlying vital rates (González & Martorell 2013), as well as easing time-series sensitivity and weakening priors (González et al. 2016). Crucially, despite these extensions, we still do not know what vital rates are most influential in the solution of inverse models of species’ life cycles. How many and which combinations of vital rate parameter uncertainty are most likely to yield identical values (*i.e.,* non-identifiability) that would limit the precision of inference available through inverse modelling? How sensitive is posterior inference to prior information of the life cycle? What influence does life history exert on inferred vital rates? Answers to these questions will increase the utility of inverse models in population ecology and strategically guide the acquisition of data to fill data gaps.

To address these questions, we assess the efficacy of solving inverse problems across simulated datasets representing a range of virtual species with distinct life history strategies, priors, unknown parameters, and dataset duration. We use this modelling approach to first evaluate which unknown vital rates are most likely to cause non-identifiability in inverse problems in demography. We then examine how imprecise or inaccurate priors propagate into errors in the estimation of unknown vital rate parameters. Next, we explore how priors are most effectively used to resolve non-identifiability based on the life history of a virtual species. We conclude by evaluating the above questions through variable length time-series, thus addressing how time-series influences the ability to solve vital rate parameters from population structure time-series data. Collectively, these questions offer insights to the potential use and misuse of inverse problems in demography informed by the type of data and general life history under investigation.

## Methods

We first detail our statistical framing of the inverse problem in population ecology, followed by a discussion of the demographic modelling framework, and then apply it to six different modelling scenarios that entertain variable dataset length and life histories. These modelling scenarios combine demographic and statistical parameterisations to test the contingencies and influence of priors on the posterior probability of unobserved vital rates. The demographic and statistical parameterisations differ in the vital rate functions of the generating model, the unknown parameters solved by the inverse modelling procedure, the sampling timing and density, and the strength and specificity of the priors.

### State Space Models and Parameterisation

To infer vital rates from time-series of population counts, we use a type of general state space model (SSM). SSMs are a broad class of statistical models that are distinguished by the inclusion of unobserved (hidden / latent) parameters in their model structure. The hierarchical SSM model partitions sub-models to estimate observed and unobserved parameters (*i.e.,* latent and process, respectively, *sensu* McClintock et al. 2020) simultaneously. The partitioning of variance between observed and unobserved parameters is achieved via a hidden process model that explicitly distinguishes processes underlying identical data patterns (*i.e.*, non-identifiability). Non-identifiability describes many-to-one associations of data to parameters, which can preclude assignment of probability under traditional statistical frameworks. For instance, if one attempts to estimate the biomass of two fish species using size data *alone*, and if two fish species have individuals of the same size, then fish of the overlapping size range cannot be uniquely linked to species and are therefore nonidentifiable. As a generalisation, when observed data have fewer dimensions than the number of unknown process parameters, models become structurally predisposed to identify single observed states consistent with multiple combinations of parameters (Wieland et al. 2021). Parameter identification defines the scope of utility of the inverse problem of population ecology and is related to a mismatch in the degrees of freedom in population structure and vital rates (Caswell 2001). For instance, a standard 3×3 Leslie (*i.e.,* age-based) matrix population model would in its simplest representation have two survival matrix elements on the sub-diagonal and one reproductive element on the right-most element of the first row, and each time-step in this age-structured population time series would have three elements. SSMs provide the framework to support inference of parameter identifiability behind these models and can help estimate the minimum conditions necessary for parameter identifiability (De Valpine 2003).

Particle filter approaches are a powerful and flexible method for solving dynamical systems using SSMs (Storvik 2002). Particle filters (also called Sequential Monte Carlo (SMC); Doucet, Freitas, & Gordon 2001) are a class of Approximate Bayesian Computation (ABC) models that optimise distance from simulated outcomes to fit parameters *in lieu* of conventional likelihoods (Toni et al. 2008). Unlike Kalman filters (1960), particle filters do not make any Gaussian assumptions about underlying parameter distributions. Particle filters have been used in multiple applications in population ecology, from density-dynamics of red kangaroo (Knape & DeValpine 2012) to environmental controls on phytoplankton (Dowd, Jones, & Parslow 2014) to grey seal metapopulation dynamics (Thomas et al. 2005). Our framework here employs a particle-filtering-based approach to infer unobserved vital rate parameters. The particle filtering approach operates through the basic concept of Importance Resampling, whereby parameter estimates are iteratively resampled as a function of their goodness of fit (Fig. 2). Particle filtering generates a filtered particle density approximating the true posterior probability of parameter values. Here, filtering refers to the process of using more recent data to update inference from the prior values in a state vector, thus *filtering* information to preceding timesteps. More detailed information about the SMC is described in the *SMC Routine* section, below.

**Figure 1.**
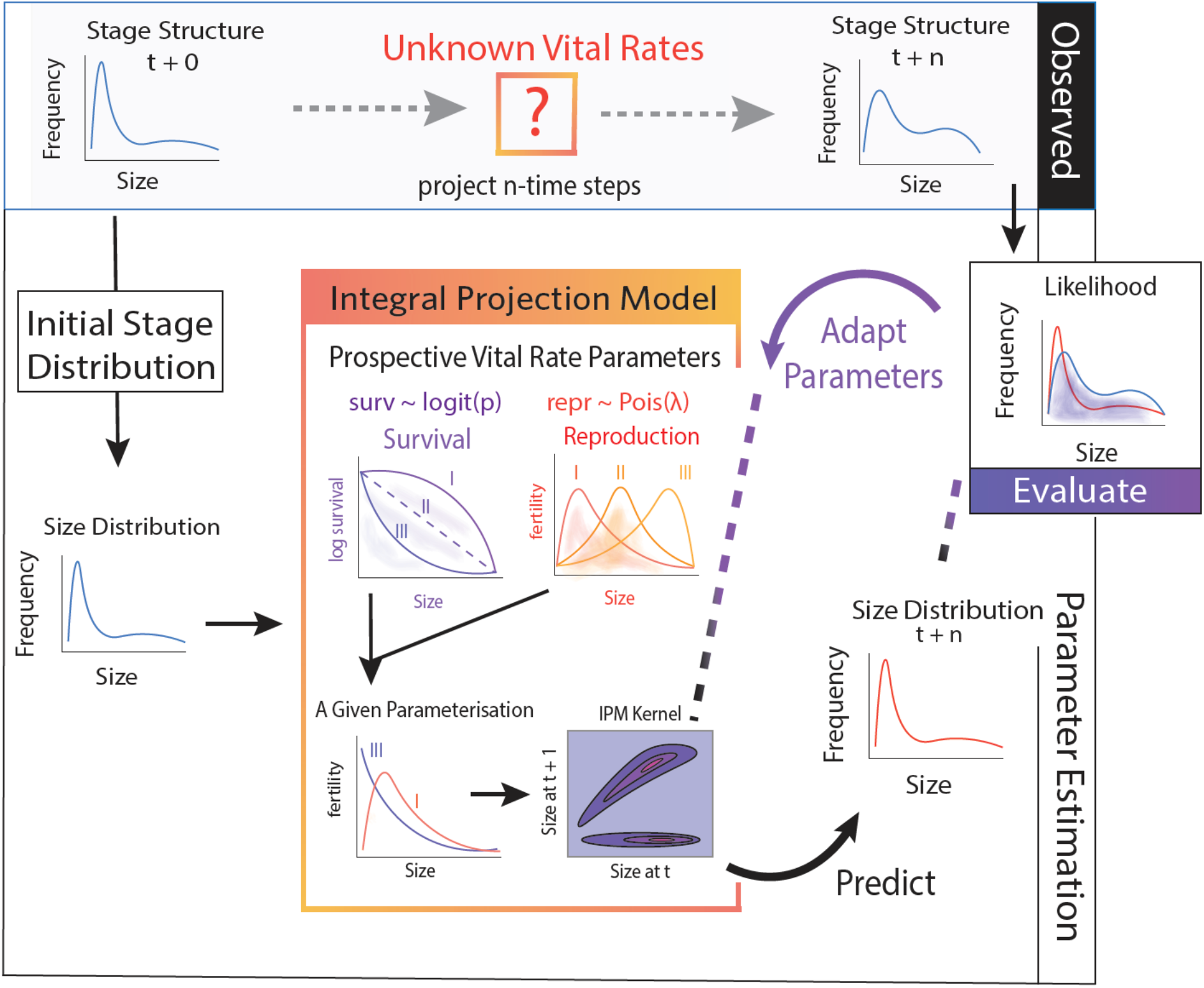
Adaptive resampling framework for solving unknown parameters of a given Integral Projection Model (IPM) using inverse models. This approach uses time series of population stage structure (*i.e.*, number of individuals of a given state value; *e.g.*, size) as the basis for the performance of an IPM. Temporal shifts in stage structure of a population over time (top row) reflect outcomes of a closed set of parameters of linear functions that control the transition of stage through an Integral Projection Model (*i.e.,* a vector function that behaves as a matrix operator). The probability mass (typical set) of parameters that can generate the observed state transition sequence may be identifiable or nonidentifiable for a specific parameter across other parameter combinations (see Figure 2 for more detail). To estimate the set of solutions for a given series of stage-transitions, one must efficiently sample from the integral of the probability mass to obtain a measure of its performance through state space. Here we use a particle filtering algorithm as the basis of our probability integral sampling (see Figure 3 for more detail). At its highest level, specific parameters are sampled that correspond with particular linear functions (top row of Integral Projection Model insert). The parameter values sampled in a given iteration are linked through an IPM (bottom row of Integral Projection Model insert) and project the initial state vector forward time (Predict Step, lower right). At each time-step performance of parameters is measured using a Shannon-Jenzen distance (operationally identical to negative log square distance). Based on performance in each preceding step, a filtered set of parameter values are retained and re-sampled (Adapt Parameter feedback step, upper right). With the iteration of the algorithm, samples maximize expectations of parameter values, converging to a time-invariant set of parameter values corresponding with comparable performance.

**Figure 2.**
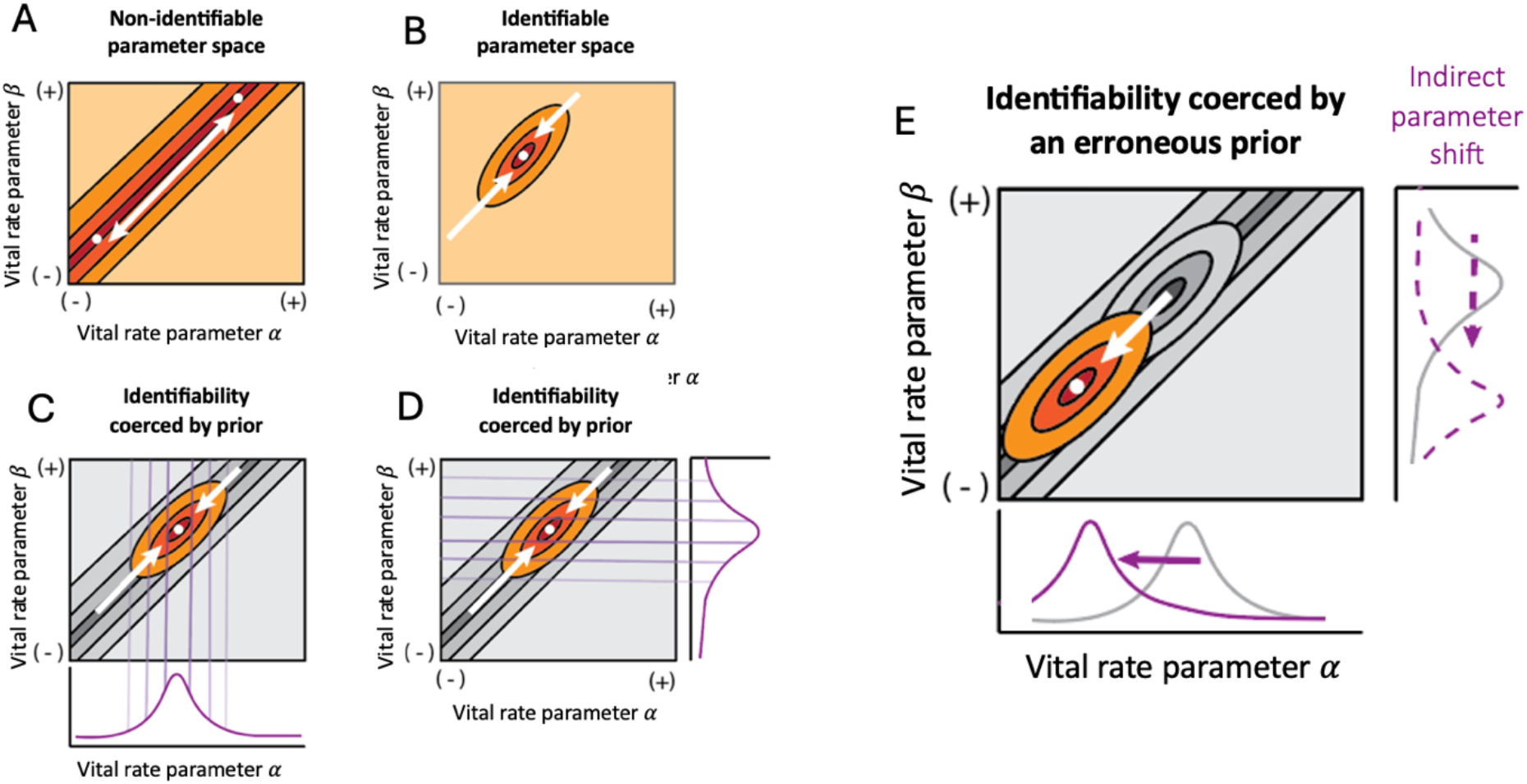
Graphic depiction of nonidentifiability in parameter space within an integral projection model (IPM). Maximum likelihood or minimum distance to the true time series as estimated through optimization methods or other time-series fit includes a set of solutions that pair values of two parameters (Panel A). In such a scenario, one can only estimate the potential parameters of the IPM to a ratio of parameter values, but not any particular parameter value (Panel A). Bayesian methods using informative priors constrain the set of potential solutions for one or more parameters such that the probability defined for one parameter can help resolve one or more additional parameters (Panels C & D). That is, informative priors (such as those illustrated in Panels C & D) can render nonidentifiable space (*e.g.,* Panel A) into identifiable space (*e.g.,* panel B) by selectively excluding portions of the nonidentifiable set. When using informative priors to resolve identifiable solutions from nonidentifiable sets in the uninformative space, modifying the priori distribution on one parameter can result in movement in the posterior density for another parameter with little bearing on accuracy (Panel E). This diagram illustrates how priors can exert strong changes parameter estimates of inverse problems. In this study, we consider whether the depth of sampling (increasing the training data) could putatively shift this distribution.

**Figure 3.**
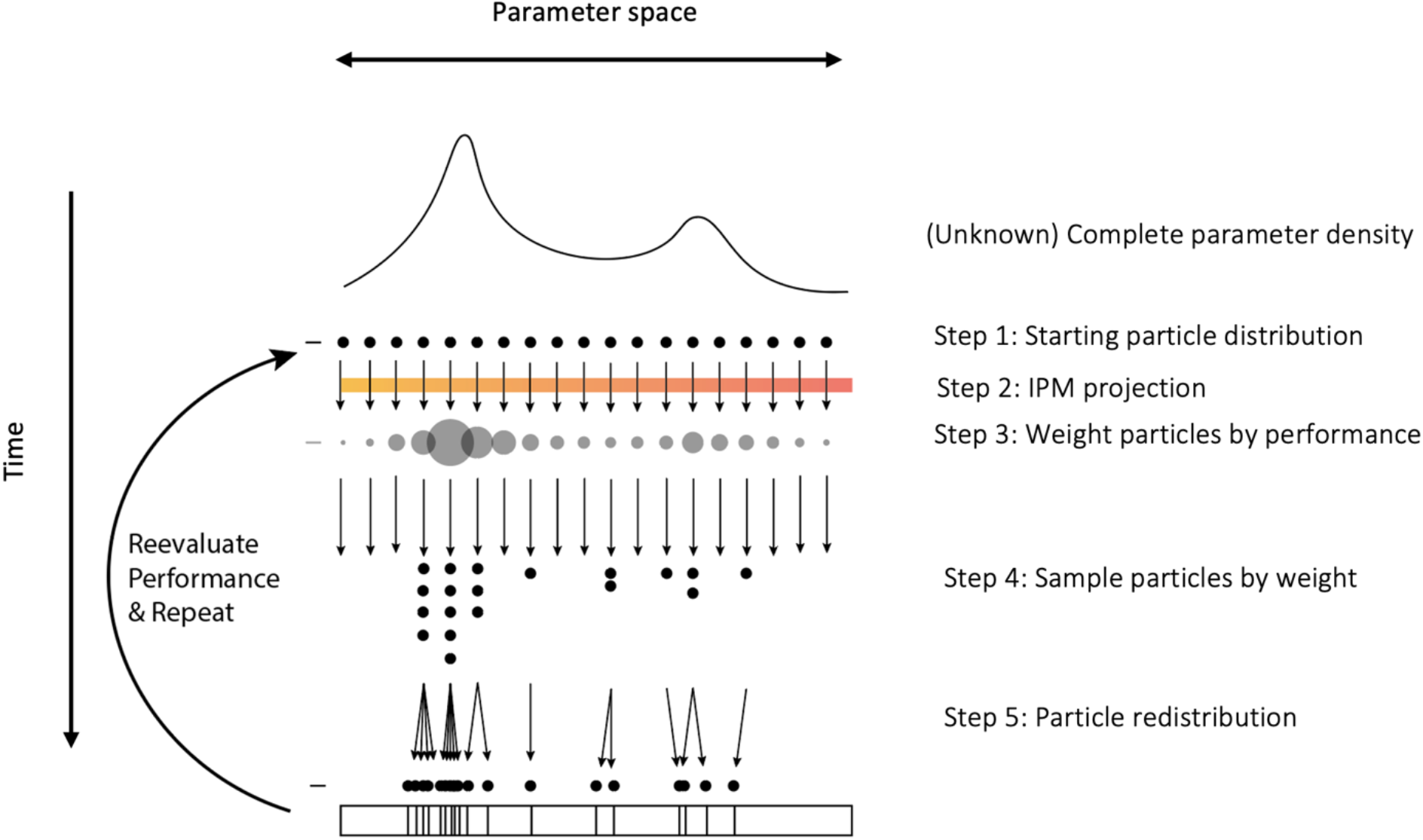
Conceptual diagram of Particle Filtering Importance Resampling used in this study to estimate unknown vital rates by sampling the integral of the probability of parameters whose performance is based on population structure time-series. The framework is based on the iteration loop: Step 1: Particles begin with a distribution based on the prior or the posterior from the preceding iteration; Step 2: Particles are used to project a population time series by parameterizing functions integrated as an IPM; Step 3: Particle performance is measured by the difference between the stage-structure projected from parameters and that of the observed time-series; Step 4: Particles are resampled weighted by performance; Step 5: Particles are perturbed on a time-dependent kernel to prevent particle degeneration.

### Integral Projection Model

Integral projection models (IPMs) are discrete-time population models where vital rates are structured by continuous state variables (*e.g.,* size or height; Easterling et al. 2000; Ellner, Childs, & Rees 2016). Here, for the purposes of tractability, we entertain a deterministic, density-independent IPM with time-homogenous vital rates, in a spatially homogeneous, closed population – *i.e.* no migration. We keep our models simple to highlight basic properties of interest; note, however, that more complex IPMs exist, which can incorporate stochasticity, density, and multiple states (Ellner & Rees 2007; Vindenes, Engen, & Sæther, 2011; Coulson 2012).

IPMs typically require fewer parameters to define the life cycle of a species than MPMs (Ramula, Rees, & Buckley 2009; Ellner et al. 2022), thus rendering identifiability more likely for IPMs than MPMs in the context of inverse modelling (González et al. 2016). The generalised linear models that are typically used in the construction of IPMs are responsible for this technical advantage with respect to parameter identification. In IPMs, one must estimate parameters of the distribution of vital rates (survival, changes in state [*growth* if the state describes size, volume, or weight], reproduction) and is not required to estimate vital rates for each discrete state, unlike in most MPMs (Caswell 2001). When regressions are compiled into an IPM, the distribution of individuals of state *z* at one time step *t* becomes mechanistically linked to the distribution of individuals of new state values (*z’*) at *t*+1 via a kernel, or linear map. In these models, *z* is a value within the compact set of possible state values ***Z***. The time-specific state structure of the population (*d_t_(z)*) is a time-explicit state distribution whose value is shaped by vital rate distribution functions (*e.g.,* survival *s(z)*, growth *g(z’,z)*, reproduction *f(z’,z)*).

In relatively simple life histories (*e.g.,* no clonality, no dormancy, no migration), IPMs are typically composed of two constituent sub-kernels: a *P*-kernel, which details changes from *z* to *z’* conditional on survival – here growth for simplicity; and the *F*-kernel, which links the size *z’* of new recruits in time *t*+1 to the reproductive efforts of parents of size *z* in time *t*. For simplicity, the *F*-kernel (*F(z’,z)*) consists of the joint distribution of size-specific fecundity and recruitment in *t*+1 as a function of recruitment state *z’* values. We use a linearized IPM, constraining the relationship between survival, growth, and fecundity to simple, linear functions:

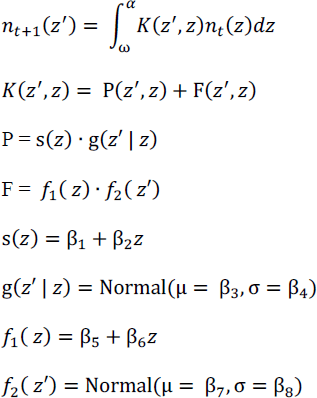

Where *s(z)* defines survival, *g(z’ | z)* defines growth, *f_1_(z’ | z)* defines fecundity, the number of offspring produced per capita, *f_2_(z’)* defines recruitment, the distribution of newborns *z’* values at *t*+1. The range of states that an individual can take is defined by the state spectrum constrained by α and ω. For simplicity again, we have fixed a constant recruitment probability that multiplies the fecundity term in the reproduction function with a constant value (but see Discussion). Logically, we also bound survival between 0 and 1, and reproduction to be non-negative.

### Simulation Routine

To infer vital rates from time-series population size and structure data, we first consider the observed state sequence (*c_1_, c_2_, … c_n_*). This sequence represents the frequency of individuals of different state values (here size) at time *t*. A time-series of state vectors constitutes the state of an initial population structure (*v_1_, v_2_, … v_n_*) transformed by the IPM’s vital rates over time. Thus, observed state sequences reflect underlying vital rate parameter combinations. The true, unobserved variables (*r1, r_2_, … r_n_*) constitute potentially unique solutions to the observed state sequence or may represent a single instance of a closed or unbounded set of parameter values consistent with the observed data. The observed state vector reflects a Markovian (*i.e.*, memoryless) process, with each time-step dependent upon both the individual vital rates and population structure in the directly preceding time-step.

To test the effect of priors and prior specificity on the accuracy and precision of posterior estimates of vital rates, we initially evaluate a simple, linear deterministic IPM. To do so, we first linearise vital rate functions and keep the IPM to the time-homogenous, density-independent case. Next, we evaluate how time-series duration, priors, and life history inform inverse solutions to IPM parameters. Finally, we generalise lessons that extend to more complex model forms, as addressed in the Discussion.

To estimate vital rate functions underlying time series, we used an SMC approach. We configured an SMC for this study following a conventional SMC framework (Doucet, Freitas & Gordon 2009; Cappé, Godsill, & Moulines 2007). We initiated the SMC by drawing independent vectors from θ = (θ_1_, θ_2_, … θ_v_), with each vector containing vital rate parameters based on a prior distribution that we specified for each vital rate (π_i_). Each vector in θ represented a candidate IPM and life history with corresponding demographic properties (*e.g.,* a high reproductive intercept suggests early reproduction and longer reproductive window, all else being equal; a low intercept and low reproductive slope indicates delayed, less intense reproduction over a shorter reproductive window; a high survival intercept and low slope correspond with life expectancy, and slow life history strategies (*sensu* Stearns 1983). Here, vectors are synonymous with particles. We define an initial population distribution structured by our state variable (***n****_1_, **n**_2_, … **n**_n_*). We then project a population forward in time following the vital rates specified by a given particle. We evaluate the performance of each particle using a distance function, measuring the goodness of fit of each particle at time-step *t*. To quantify the distance between the observed and simulated stage distributions, we use the Jensen-Shannon Distance (JSD). JSD is an unbiased, symmetric measure of distance between probability distributions (Lin 1991).

To iteratively improve the efficiency of the search algorithm, we employ computational heuristics (Olafson 2006), which shortcut raw computation using approximation rules. Optimisation heuristics describe a class of procedures that isolate a nearly exact solution to optimisation problems without the computational cost of evaluating the exact mathematical solution (Glover & Greenberg 1989). Because heuristics cannot guarantee a correct solution on a single run, they benefit from ensemble approaches and validation with complementary methods (Bianchi et al. 2006). We use two heuristics at the resampling step, between the proposal and test distribution. Low-performing particles are not resampled based on a prohibition threshold that we prescribe, restricting the resampling step to a parameter space of higher performance. This heuristic, similar to a tabu search heuristic (*sensu* Glover 1990), improves the efficiency of convergence in the optimisation algorithm. To further improve the efficiency of the search algorithm, we also use a performance weighting parameter whose regulation operates similarly to simulated annealing. The performance weighting parameter is initially set at a high temperature, discounting the performance measure’s influence on resampling, encouraging resampling across the initial parameter space. As the performance weighting temperature cools with increasing resampling, only the highest value particles are resampled.

### Sequential Monte Carlo Routine

We used a sequential Monte Carlo routine to estimate parameters of the vital rates comprising an IPM-generated stage-structure time sequence. We initiated a population vector (***i***) at the first-time step (*t* = 0). Particles were drawn from prior distributions for each vital rate parameter (θ*_i_*∼π_θ_). We simulated a stage structure at time *t*, for each particle (*x_it_*∼*M*(θ; *i*)). We calculated the distance between the time-specific stage structure between *t*-1 and *t*. The performance of each particle was calculated by the distance to the observed stage-structure value and normalised with respect to the particle number (1/*N* = *w_i_*). In discrete intervals across the domain of the particles, we retained the highest performing particle, and assigned it to the mid-point parameter value of that interval (*w_i_*). Operationally, this step marginally maximises the parameter-specific performance across all other parameter values.

Following from the above, we iterated the following steps described in Figure 1. We reset a threshold that determined the minimum weight of particles that could be resampled. We updated the scaling parameter of particle weights (increasingly biasing higher sampling in areas of higher parameter performance). Next, we resampled with replacement of each particle (*θ*) with the normalised weighting probabilities (*wi*). We perturbed the resampled particles (θ*_i_*∼*N*(θ, 0.001)). We then calculated the performance weighting (distance) of the particles. We discarded particles whose distance exceeded that of the threshold defined in the first step of this iteration procedure. We then normalised the performance weightings, and returned to the first step in this iteration procedure.

### Demographic Scenarios

To the best of our knowledge, all empirical treatments of the inverse problem in population ecology to date have modelled existing time-series data. These efforts have demonstrated the ability to indirectly infer unobserved vital rates on a handful of relatively long-term datasets on a limited set of life history strategies (*e.g.,* Tavecchia et al. 2009; Caswell & Twombly 2012; Gross, Craig, & Hutchinson 2012; see notable exception with the ten synthetic species in González & Martorell 2013). To strengthen our understanding of how life history strategies influence the estimation of vital rates using population time series, we compared the accuracy of inverse modelling for five life histories that vary in their generation times from from five to 30 time-steps. Generation times provide a single-value demographic measure that correlates with a broad set of demographic traits (Gaillard et al. 2005) and affords a proxy for variation along the the fast-slow continuum (Stearns 1989). Each life history strategy was described by three vital rate functions (survival, growth,and reproduction) and eight vital rate parameters (Fig. 1). Life history strategies were constrained such that intrinsic population growth rates (λ) were bound to 1 ± 0.05 S.D., and which varied in the intercept and slope parameters for either reproduction or growth functions. For survival, intercept was bound between 0 and 1 to adhere to the biological and mathematical limits of this process. For growth, intercept and slope were bound between 10 and -10, representing extremely rapid growth and shrinkage, respectively. We constrained survival and reproduction functions to be monotonic, such that both vital rates do not change their relationship to growth (*e.g.*, increasing then decreasing). Net reproductive rates were limited ot non-zero, non-negative values. We constrained the resulting IPMs to vital rate parameter values that did not result in infinite or incalculable generation times nor net reproductive rates, or IPMs that, after discretisation (Easterling et al. 2000), would result in nonreducible or negative matrices.

We accounted for differences in the stationarity and transient (*i.e.*, short-term) dynamics of populations (Stott et al. 2011). To do so, we simulated population time series across three scenarios of initial population structure: stable st/age distribution (represented by projections to 1,000 time-steps); a stage structure where the population is initially compacted into the lowest value structure (*i.e.,* a vector with value = 1 in the first state position, trailed by 0s); and the corollary stage structure where the population is concentrated into the largest state value (*i.e.,* a vector with value = 0 in all positions by the last one, which has a 1).

### Statistical Scenarios

To evaluate the importance of priors in posterior Bayesian inference of vital rates, we quantified the sensitivity of posterior estimates of vital rate parameters to priors. Here, specifically, we expected that the effect of prior precision and accuracy on posterior estimates should be affected by time-series length because more information about vital rates is expressed in population structure with time. To account for these parameters, we evaluated a range of time-series and parameter estimation scenarios.

### Parameterisation Scenarios

To evaluate the effectiveness with which parameters are recovered in our framework, we quantified all scenarios where two parameters were simultaneously estimated successfully. For instance, we tested the ability of our framework to estimate both the slope and intercept of the survival function, and then the ability to estimate the slopes of the survival and reproductive functions. In doing so, we assumed knowledge of all vital rate parameters barring two out of the total of eight (Fig. 1), and used population structure to estimate the unknowns and permute this estimation procedure across all parameter combinations. We used two tolerance thresholds to estimate a binary status of whether a posterior is sufficiently close to the true value to be considered identifiable for the purposes of summarising data.

Taking this dual threshold approach allowed us to investigate two key aspects regarding the performance of our framework: the first threshold addresses spread (S.D. = 0.1), while the second threshold addresses accuracy (maximum distance to the true value of 0.1). Wide posteriors indicative of nonidentifiability may have central tendencies that fall close to the true value, which is why we used both variance and median as a joint criteria. Unless noted elsewhere, the tolerance criteria were applied to both unknown parameters for the purposes of data summaries. As the IPM used here includes eight vital rate parameters, the maximum number of identifiable combinations was bound between one and eight (the focal parameter plus one of eight remaining values).

## Results

IPM vital rate parameters were correctly estimated using population structure in a synthetic environment. We used a particle-filter based inverse modelling procedure to identify parameters in a pairwise fashion, whereby two of eight parameters defining three vital rate functions (survival, growth, and reproduction) were unknown and solved using time-series of population structure of variable length and life history characteristics. This procedure allowed us to do a complete parameter estimation sweep within the computational constraints of the models and generate nonidentifiable scenarios in a simple case where multiple solutions exist. The vital rate parameters accurately estimated using population structure were a subset of the total pairwise combinations and varied by parameter type (*e.g.*, intercept, slope) and vital rate (*i.e.,* survival, growth, reproduction). Certain parameter combinations were persistently nonidentifiable at short- and medium-term time steps, while others were consistently discernable. For instance, combinations of reproduction parameters and standard deviation of growth, or growth slope and survival slope, were not discernable. Moreover, contrary to our expectations, life history and time-series duration explained a small proportion of the variance of parameter identifiability. Importantly, when more information about vital rates was supplied using priors, the strength of indirect inference on remaining unknowns improved (Fig. 2).

Life history strategy, as classified by generation time, generally did not affect parameter identification across the full set of vital rates (Fig. 4). In seven out of eight pairwise parameter estimation models, the 95% credibility intervals for the slope parameters overlap with zero, implying no effect. The predictive power of generation times, ranging from five to 30 time steps, on parameter vital rate estimation was null for both transient and asymptotic phases for all parameters except for the slope of the growth vital rate regression. The identifiability of vital rates was conditional on stationarity for survival parameters (intercept = 4.8 [2.78; 6.87] (stationary); 3.69 [-0.94; 8.16] (transient); slope = 5.37 [2.35; 8.63] (stationary); 3.75 [-0.36; 7.81] (transient)) and reproductive slope (2 [2;2] (stationary); 1.09 [-0.90; 3.20]) (transient)). Identifiability did not differ in the significance or sign for the growth parameter estimates. The effect of generation time on growth slope was comparable between transient and stationary models (0.816; 0.182, respectively), but the estimates were marginally insignificant for the transient model (0.186 [0.03; 0.34])).

**Figure 4.**
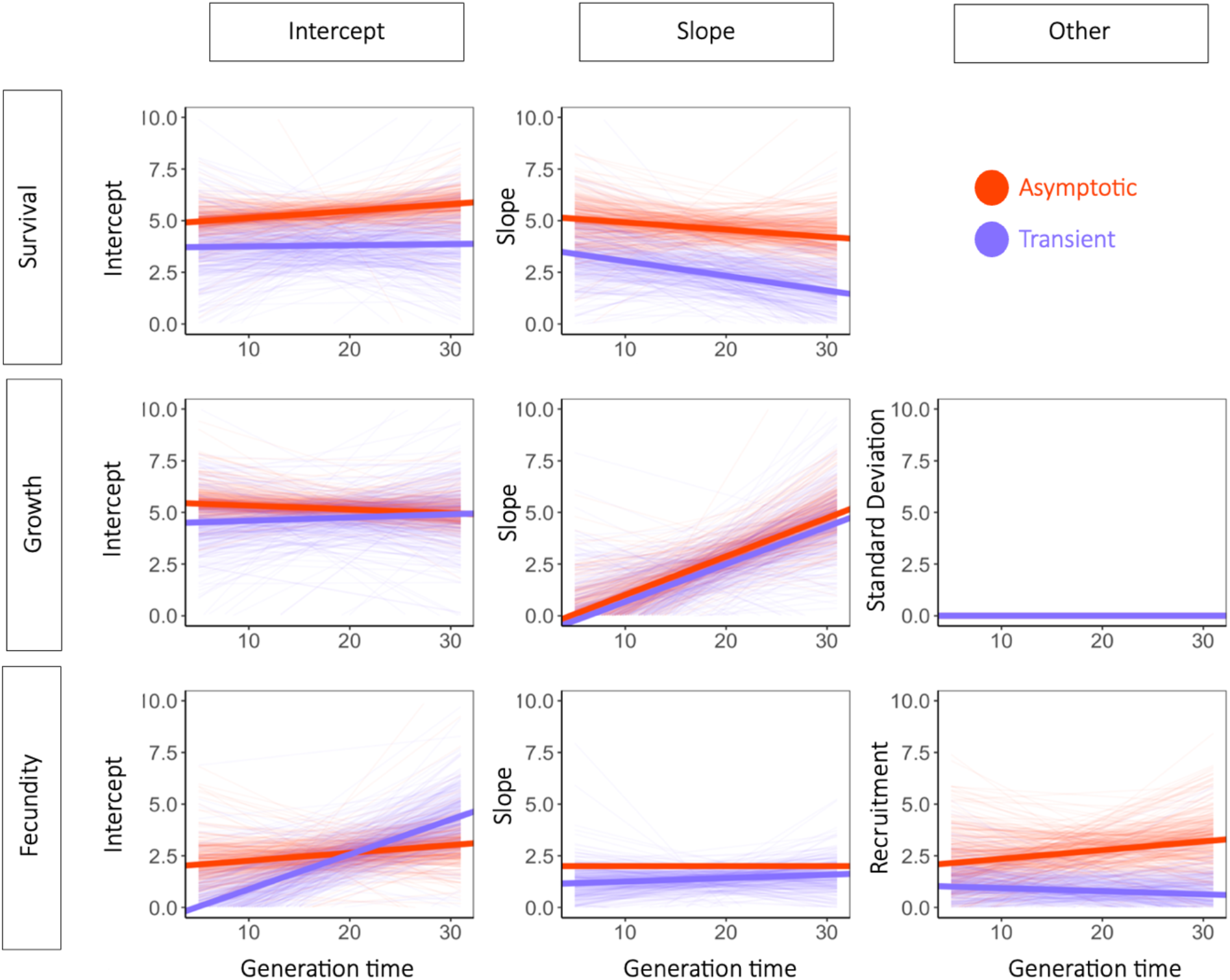
Generation time is a poor predictor of parameter identifiability. Effects of generation time on the number of identifiable vital rates in a pairwise knockout. All combinations of pairwise parameters were solved by fitting a five timestep population time series. Where posterior estimates of vital rate parameters were within 0.01 of the true parameter value, we considered them identifiable. Each pair of vital rates was solved across all seven pairwise unknowns, thus the maximum number of identifiable outcomes for that parameter in a given scenario was seven. Generation times ranged from 5-30 time steps. Asymptotic lines (purple) reflect solving the inverse problem when the initial population structure was the stable stage distribution for the IPM analysed; Transient lines (orange) reflect scnearios in which the population was perturbed from its stable stage distribution at is furthest extremes. The slopes of these correlations are not significant. Dark lines reflect the mean response. The semi-transparent lines show 100 draws from the posterior probability.

Prior strength in vital rate priors improved the ability to correctly identify vital rate parameters for growth. Other vital rates showed insignificant but consistent trends toward the same effect of prior knowledge (Fig. 5). Growth intercept (-1.34 ± 0.63) and growth slope (-1.55 ± 0.69) improved parameter identification with stronger priors (*e.g.,* SD < 0.25). Coefficients were universally negative, consistent with the directional trend of higher identification with stronger priors, but were insignificant likely owing to low statistical power. Parameter identifiability varied amongst transient and asymptotic scenarios for survival (intercept = 3.24 [1.37; 5.28] (asymptotic); 6.35 [5.19; 7.47] (transient)). Identifiability did not dramatically or systematically differ across survival (intercept = 4.78 [3.34; 6.26]; slope = 4.14 [2.34; 5.99]), growth (intercept = 5.08 [3.70; 6.57]; slope = 3.38 [1.82; 5.01]), and reproduction (intercept = 3.19 [1.80; 4.64]; slope = 2.65 [1.35; 3.99]).

**Figure 5.**
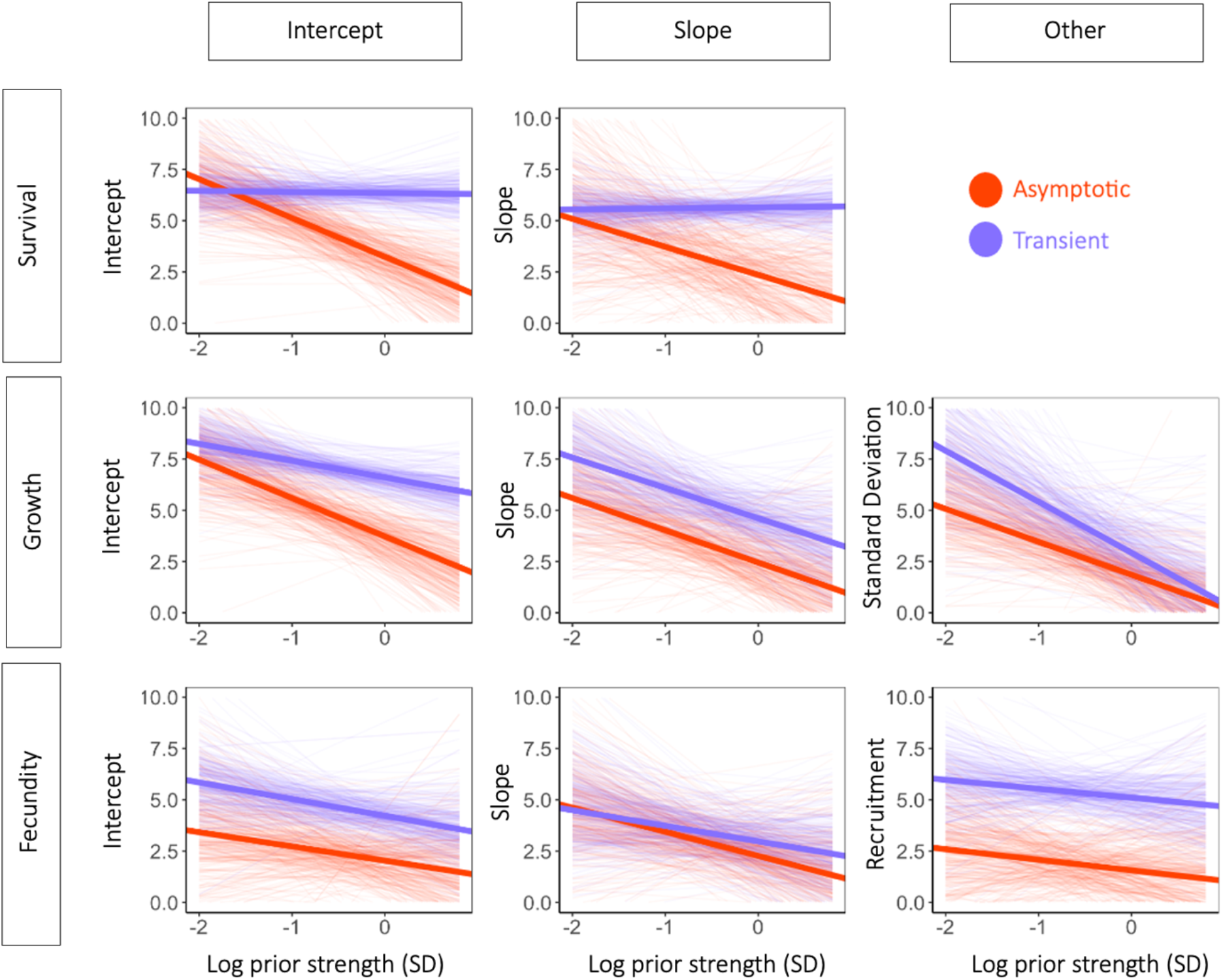
The strength of priors on unknown parameters has a critical effect on the ability to identify unknown parameters without informative priors. Effects of standard deviation on a normal prior applied to the compensating parameter on the number of identifiable vital rates in a pairwise knockout. All combinations of pairwise parameters were solved by fitting a five timestep population time series. Where posterior estimates of vital rate parameters were within 0.01 of the true parameter value, we considered them identifiable. Each pair of vital rates was solved across all seven scenarios, so the maximum number of identifiable outcomes for that parameter in a given scenario was seven. Prior strength ranged from 5 to 0.01 and is plotted on a log scale. Where the stage structure is concentrated to late stages, this is shown in purple and labelled transient, because sampling of the population time series reflects the transient envelope. Where the initiating stage structure is the stationary stage structure of the population (i.e., time-homogenous / asymptotic), this is shown in orange and labelled asymptotic.

Contrary to our expectation, time-series duration did not improve the ability to identify vital rate parameters in the asymptotic or transient phases of estimation (Fig. 6. In all eight pairwise parameter estimation models, the 95% credibility intervals for the slope parameters overlap with zero (S1). However, there was a difference between the number of vital rate parameters successfully estimated in the asymptotic *vs.* the transient phase, with higher estimation likelihoods for the asymptotic conditions (p < 0.05 in 63% of the parameters; mean difference = 1.96). Higher rates of identifiability occurred in the vital rates for survival (intercept = 4.7 +/-1.64 sd; slope = 3.7, +/-1.77 sd) and growth (intercept = 5.4 +/-0.51; 4 +/-0.47) compared to reproduction (intercept = 2.8 +/-0.78sd; slope = 0.90 +/-0.10 sd). Growth and survival comparisons with reproductive parameters were roughly equivalent (mean = 2.85 [2.50; 3.19 95% CI]; 2.35 [1.63; 3.07], respectively).

**Figure 6.**
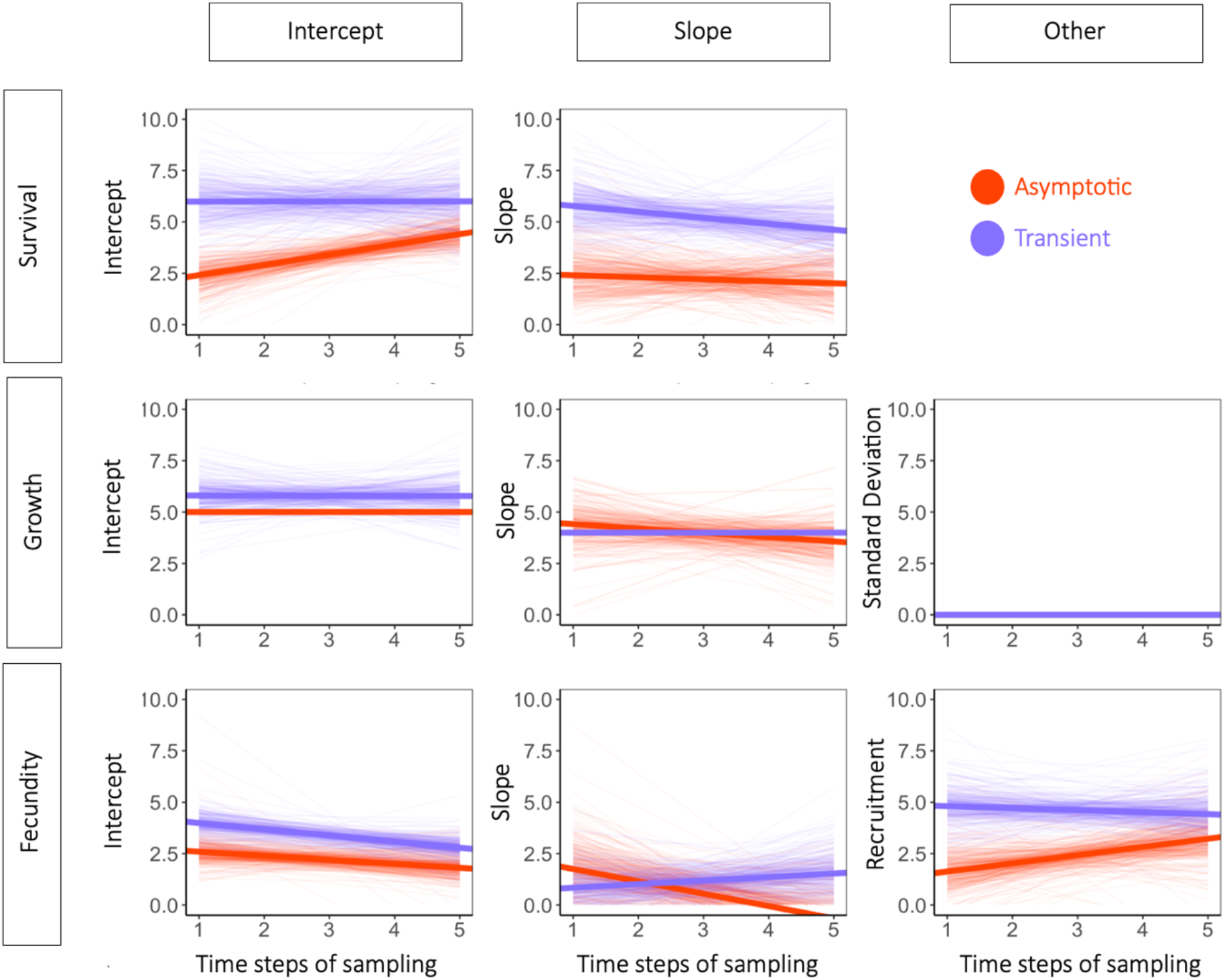
Time series duration does not improve parameter identification in inverse models. Effects of sampling duration on the number of identifiable vital rates in a pairwise knockout. All combinations of pairwise parameters were solved by fitting a five timestep population time series. Where posterior estimates of vital rate parameters were within 0.01 of the true parameter value, we considered them identifiable. Each pair of vital rates was solved across all seven scenarios, so the maximum number of identifiable outcomes for that parameter in a given scenario was seven. Sampling duration ranged from one to five time steps. Where the stage structure is concentrated to late stages, this is shown in purple and labelled transient, because sampling of the population time series reflects the transient envelope. Where the initiating stage structure is the stationary stage structure of the population (*i.e.*, time-homogenous / asymptotic), this is shown in orange and labelled asymptotic. None of the slopes were significant.

## Discussion

Study duration remains one of the most persistent limitations in the construction and utility of population models in pure and applied ecology (Ward et al. 2014; Doak, Gross, & Morris 2005; Keyfitz 1981). The lack of long-term demographic data reflects the cost of long-term monitoring (Treurnicht et al. 2016; Pollock et al. 2002). While innovation in field technology make tracking of individuals and their populations more accessible and cost-effective (*e.g.*, UAVs (Olsoy et al. 2024; Stone & Davis 2023); passive song-meters (Kahl et al. 2021); highly-resolved GPS (Williams et al. 2014; Willoughby 2017); eDNA (Yates, Fraser, & Derry 2019)), large-scale shortfalls in the availability of demographic data for species will likely persist (Buckland et al. 2023). In the face of foreseeable demographic data deficits, inverse modelling has emerged as a promising approach to extend demographic inference without demographically explicit data (Bruijning, Jongejans, & Turcotte 2019; Evans et al. 2016; González et al. 2016). Research treatment of this subject to date has been limited to species-specific applications (*e.g.,* Félix-Burruel et al. 2021 (saguaro cactus, *Carnegiea gigantea*); Ghosh et al. 2012 (tulip, *Liriodendron tulipifer*); White et al. 2016 (rockfish, *Sebastes mystinus*)), irrespective of short-time-series, and largely indifferent to how model structure interacts with demography to influence estimates. Here, using an inverse integral projection model (IPM) approach (González et al. 2016) to estimate vital rates using population structure in a Bayesian framework, we show that longer time-series duration does not substantially improve the ability to solve IPMs based on stage structure alone, that solving IPM vital rate parameters is insensitive to life history differences, and that comparatively few parameters can be solved at any given time without relying on strong priors.

Increased time-series duration has been regarded as an essential requirement for improving inverse vital rate parameter estimation (González et al. 2016, De Valpine & Hastings 2002; Kendall et al. 1999). The logic here is that longer time series may reduce error in inverse modelling in at least three ways: (1) sampling duration allows partitioning of observation error from the generating process signal (Newman et al. 2023; Deitze 2017); (2) longer-term data provide more complete representation of fluctuations in the cohort cycles of a population (Botzford et al. 2019; Hastings et al. 2018); and (3) longer-term data are more likely to capture the influence of the environment on vital rate variation (Ellner, Seifu, & Smith 2002; Kendall et al. 1999). Additionally, longer-term data are expected to generate more departure between candidate vital rate values with time, because small rate differences compound geometrically (Caswell 2002). In other words, population size differences generated by small differences in parameter difference (e.g., reproductive rate) are measurably clearer after, say, ten time-steps than they are after just one or two. However, our results show that the benefits of improved precision of vital rate estimates inferred from population time series are not associated with information contained in the stage structure of the examined population. Rather, the benefits of longer time series to inverse inference may be grounded in the time-series-moderating effect of observation error, in density regulation, or other factors. This finding is particularly surprising when joined with the observation that inverse modelling is insensitive to generation time, based on the principle that information linking vital rates to time series is mediated by population structure. Our results suggest that more work is needed to better understand the key drivers of sensitivities in inverse modelling.

In the simple, deterministic case where there is no observation error and the population structure is non-stationary, one might expect identifiability of vital rate parameter estimates to improve over the duration of structural transients (Koons et al. 2005). The connection between the transient envelope, which defines all the possible growth/decline responses that a structured population can exhibit in response to an uneven disturbance to its stage structure (Caswell 2011), and parameter estimation follows from two related ideas. First, longer time series affords the ability to test how compounding differences in vital rates reflect in the age structure (Ellner et al. 2002). Second, the transition from the time-varying to time-invarying structure limits the extent to which time series translates into information on vital rates (González et al. 2016; Caswell 2002). Once the stage structure recovers stationarity, one does not expect a time-dependent effect to persist on estimates or identifiability because of the time-invariance in population size (Wachter 1986; Tuljapurkar 1982; Caswell 2002). Here, however, we found that time effects were indiscernible early in the time-series. This difference could be due to a combination of long-transients (*sensu* Hastings 2016) that exceed the sampling window in the transient samples and the expected time-homogeneity in the asymptotic samples. Or the population growth component of the model fitting could play a role of the time-dependent sampling model improvement. Indeed, while the information from population structure data becomes invariant outside of the transient envelope (Capdevila et al. 2020), solutions could improve if putative vital rate combinations have within-envelope congruence, but out-of-envelope variance.

Slower life history strategies have longer transient envelopes (Capdevila et al. 2022; Gamelon et al. 2014). When a population structure differs from a the stable stage distribution governed by its vital rates, the population structure will oscillate with a period proportionate to generation time and the damping ratio as it converges toward its stable stage distribution if stable conditions persist (Caswell 2001; Jiang et al. 2023). The speed with which a population recovers its time-invariance stationary state is proportionate to the subdominant eigenvalue of its population model (λ_’_; Caswell 2001). In a density-independent state, damped oscillations in population structure decrease with rate *e*^−t^/^τ^, where τ is damping time, defined as:

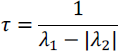

From the Lotka renewal equations, we can see that λ_1_ and λ_2_ are influenced by generation time (Wachter 1986):

Where generation time we define as:

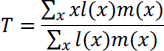

And if reproductive dispersion we define as:

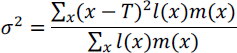

We get:

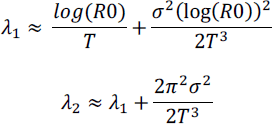

Fixing net reproductive rate, we see that reproductive dispersion will increase, λ_1_ will decrease, λ_2_ will decrease, and (λ_1_ − |λ_2_|) will decrease, in turn increasing damping time. This gives the basis for longer generation times leading to longer damping times conditional on holding net reproductive rate constant. Even if damping times are identical, longer generation times resulting in longer transients. Damping time scales allometrically with generation time based on empirical data (Jiang et al. 2022). Consequently, one might expect that longer time series are necessary to accurately identify the characteristic structural cycle for long-lived species than for short-lived species. However, we did not observe a consistent relationship between generation time and the number of parameters nor the vital rates that could be correctly identified. Ultimately, a more diverse range of life history strategies and a wider sampling window could result in patterns that are not revealed at the resolution of our study. Examples of the outer boundaries on for which structured population data exist include the Asian elephant (*Elephas maximus*), where generation times reach 20-25 years (Jackson et al. 2019), or the griffon vulture (*Gyps fulvus*), whose generation time is 1.7 (Legendre et al. 1999). See also the breadth of life histories and transients evaluated in Capdevila et al. 2022.

Past work achieved success in the inverse estimation of IPM vital rate parameters using a 100-year time series (González et al. 2016). However, the same study failed when using five-year time series, with a boundary approaching 10 time-steps and without the use of informative priors. This boundary applied to broad sweep parameterisation of the entire IPM, within the constraints of the family of functions that were prescribed *a priori*. The priors used for estimation in the shorter time series constrained the potential values on what was inversely inferred, highlighting the influential role of priors in guiding inference in shorter time series durations (González et al. 2016). Our work validates previous findings concerning the challenges of identifying multiple parameters without priors and without long-term time series (Bruijning, Jongejans, & Turcotte 2019; Caswell 2002). Our results suggest that the challenge of identifiability exists at the level of estimating as few as two parameters and does not itself resolve simply as a matter of time-series duration, whether within or outside of the transient envelope of the population of interest. Our work shows how choices of priors and *a priori* delimitation of vital rate families are critical choices for obtaining vital rate estimates from short-term time series. The ongoing growth of demographic studies (Salguero-Gómez et al. 2015, 2016; Levin et al. 2022), integrative modelling (Schaub & Kéry 2021; Zipkin, Inouye, & Beissinger 2019), and phylogenic resolution (Freckelton, Harvey, & Pagel 2002) that imputing these values might together afford an effective way to fill demographic gaps (Conde et al. 2019; James et al. 2020; Bakewell et al. 2020; Penerone et al. 2014).

Our findings reveal important features about the drivers of identifiability in inverse demographic models. Non-identifiability emerges from the interplay of compensatory dynamics amongst constituent vital rate functions in the sub-kernels of the IPM. The choice of the functional family of vital rates in an IPM can regulate the potential for multiple solutions to its vital rate parameters (Manly & Seyb 1989).

Variation in a single vital rate function and its (non)linearity may be as important, if not more important, than information introduced through priors. Changing a vital rate function for, say, reproduction from a linear map to growth to an exponential or logistic family can determine the difference in vital rate identification (not shown). Improvements in parameter estimation and identifiability over time are not driven equivalently by population growth rate, population structure, intrinsic and extrinsic variance in vital rates, density regulation, and observation error (Wood 1994; Manly 1990). While some of these factors are known to improve with longer time series (*e.g.,* sampling of the environment or observation error estimates), intrinsic factors to the models may not matter, particularly outside of the stochastic context. If model structure is not strictly prescribed and if priors are not used, then identifiable vital rates may be limited regardless of time series duration. Vital rate combinations appear to be motivated by model structure more than any particular life history, consistent with past work (Manly & Seyb 1989).

The assumptions that go into the functional form of parameters (*e.g.,* defining fecundity parameters in a linear, exponential, or logistic equation) can be determinative in whether one resolves an accurate IPM reconstruction.

Our work suggests high residual variance in the degree of identifiability that is unattributed to generation time, priors, and time-series duration. A more comprehensive treatment is merited to interrogate what life history traits, beyond generation time, might contribute to challenges in identifiability. Indeed, while generation time is a key predictor of life history strategies (Gaillard et al. 2005), the frequency of reproduction explains almost as much variation and is orthogonal to generation time (Salguero-Gómez et al. 2016; Paniw, Ozgul, & Salguero-Gómez 2018). One fruitful direction might evaluate how components of variability – directional improvement or decline in vital rates or variance structure amongst vital rates – contribute to nonidentifiability. In this context, we note that our work focused on the structural component of time-series, and not population growth. We are not aware of work to date that has partitioned the contribution to identification delivered by each of these components.

Two enduring issues that constrain the use of inverse models in demography are the lack of predictive theory concerning their usefulness under different demographic unknowns (*e.g.,* specific combinations of survival versus reproductive uncertainties), and the computational limitations that constrain nonparametric methods. The inability to anticipate the usefulness of such models *a priori* remains a barrier to their adoption and broader use in demographic studies. There is a need for guidelines grounded in life history theory for the times and places where inverse modelling is a worthwhile effort to resolve unobservable vital rates (Riecke et al. 2019; Schaub & Kéry 2021). Today, we still lack information about the relative influence of model structure, priors, and demographic traits on the identifiability of vital rates. Here, we offer an initial treatment, not a final word toward that end. The complexity of algorithms and computation time persists as the other key barrier to the adoption and wider use of inverse models in demography. Currently, inverse models remain the domain of a relatively narrow set of specialists with access to High Performance Computing and statistical fluency well beyond most demographic models. In select cases, inverse models are employed in unstructured or reduced contexts to reduce the problem to a more solvable form (*e.g.,* Ellner, Seifu, & Smith 2002; Wood 1994). There is an unmet opportunity to broaden the scope of inverse modelling with computational frameworks for solving inverse problems based on prescribing biological data, priors, and model structure. Two promising opportunities include the use of neural ordinary differential equations, which are well suited to solve inverse modelling problems (Bonnaffé, Sheldon, & Coulson 2021), and the use of parametric estimators as proxies for non-parametric tests (Glennie et al. 2023).

## Acknowledgements

We thank E. González for input in early stages of this work, as well as S. Tuljapurkar, C. Horvitz, and B. Erik-Sæther for feedback on earlier versions of this manuscript.

## Author Contribution Statement

CB and RSG conceived the ideas and designed methodology; CB wrote the simulation and modelling script; CB analysed the data; CB led the writing of the manuscript. All authors contributed critically to the drafts and gave final approval for publication.

